# Strain handling exposes IS-mediated recurrent genome reduction at a dynamic accessory locus in *Komagataeibacter rhaeticus*

**DOI:** 10.64898/2026.10.08.757553

**Authors:** Davide Bersanetti, Victor M. Gonzalez Ramos, Senni Hurttala, Anupama Prakash, Miia R. Mäkelä, Tom Ellis, Rahul Mangayil

**Affiliations:** Department of Bioproducts and Biosystems, School of Chemical Engineering, Aalto University, P.O. Box 16100, FI-02150 Espoo, Finland; Department of Bioengineering, Imperial College London, London, SW7 2AZ, UK; Imperial Centre for Engineering Biology, Imperial College London, London, SW7 2AZ, UK

**Keywords:** *Komagataeibacter*, Genome instability, Genome reduction, Mobile genetic elements, Bacterial cellulose, Phylogenomics

## Abstract

Bacterial genomes are dynamic and harbour accessory regions that can be gained, lost, or rearranged in response to selective pressures. Members of the genus *Komagataeibacter* are used for bacterial cellulose production and researched as platforms for engineered living materials. However, their genomes are rich in mobile genetic elements (MGEs), including insertion sequences (ISs), genomic islands and prophage remnants, which drive genome rearrangements under agitated cultivation conditions. Whether this genome architecture contributes to genomic changes during routine strain maintenance and distribution remains poorly understood. Comparative genome sequencing of independent *Komagataeibacter rhaeticus* iGEM stocks revealed recurrent loss of a 210 kbp chromosomal region flanked by homologous IS5 sequences. This deletion was identified in a stock received as an agar stab and reproduced from an intact-genome population subjected to simulated agar-stab handling. Phylogenomic analysis of the affected locus revealed gene-tree/species-tree discordance, heterogeneous GC content and multiple ISs, supporting its characterization as a dynamic accessory region. Populations carrying the larger genome reduction showed improved growth and cellulose production relative to the intact genome reference, whereas populations carrying a smaller nested deletion showed reduced performance. These findings demonstrate that routine strain handling is not necessarily genomically neutral and can expose structural genomic variabilities that generate population-level genomic heterogeneity with phenotypic consequences. For bacteria with MGE-rich genomes, handling history may represent an underappreciated source of genomic and phenotypic variation between laboratories. More broadly, identifying unstable accessory regions may highlight genomic content that is dispensable under laboratory conditions, providing candidates for genome streamlining and strain improvement.

**Importance:** Bacteria are commonly shared between laboratories in small tubes filled with agar gel, a practice known as agar stab. This study shows that this common practice can trigger the unexpected loss of a large genomic region in Komagataeibacter rhaeticus. Identifying such genome instability is important as this bacterium is increasingly used for industrial bacterial cellulose production. Bacteria carrying the larger deletion grew faster and produced more cellulose, while those with a partial deletion performed worse. Our findings highlight how bacteria shared between laboratories may carry overlooked genetic differences that can contribute to phenotypic differences among research groups and provide potential targets for future genome reduction engineering to improve growth and cellulose production.

## 1. Background

Bacterial genomes are dynamic entities in which core genetic functions coexist with accessory sequences that can be gained, lost, rearranged, or amplified in response to environmental and selective pressures. Mobile genetic elements (MGEs), including insertion sequences (ISs), genomic islands, prophages and plasmids, are major drivers of this genomic plasticity, and their instability can generate substantial genetic heterogeneity within otherwise clonal populations (1). Members of the genus *Komagataeibacter* are widely used for bacterial cellulose (BC) production and have increasingly been developed as platforms for engineered living materials research (2). Specifically, *K. rhaeticus* has become an established chassis for engineering cellulose-producing cells and cellulose-based materials. (3–6). A fundamental challenge in using *Komagataeibacter* as a reproducible platform is the inherent genome plasticity (7). Comparative genomic studies have identified diverse MGEs across this genus that contribute substantially to its accessory genomic content (8,9) and provide substrates for genome rearrangement, gene disruption and large-scale sequence loss (10). This can be a particular liability in biotechnological settings, where strains are expected to retain stable phenotypes during routine propagation and storage (11,12).

Genome instability in *Komagataeibacter* has been most clearly documented under agitated cultivation, where phenotypic variants with impaired BC production arise (13) through mutations and insertion events affecting the cellulose biosynthetic loci, often involving IS elements (14,15). Our previous work similarly identified multiple single-nucleotide variants and insertions/deletions affecting BC-associated loci following agitation, accompanied by the emergence of distinct subpopulations (16).

However, genome instability is not restricted to conditions that directly alter cellular physiology. Long-term storage and active laboratory passaging have all been associated with genomic rearrangements, point mutations and large-scale IS-mediated structural changes. (17–19). Yet, whether comparable large-scale structural changes can be triggered by short-term passive storage, typical of routine strain distribution between laboratories, remains poorly understood. This question is particularly relevant to *Komagataeibacter*, given its extensive MGE content (9).

Here, we investigated the genomic basis of an unexpected phenotypic difference between independently maintained *K. rhaeticus* iGEM glycerol stocks (also referred to as stocks) using comparative and long-read sequencing. We identified a large-scale deletion associated with an observed phenotypic difference and subsequently reproduced it under simulated agar-stab conditions using a parental strain with an intact genome. We then characterised the genomic architecture of the affected locus and compared growth kinetics and BC production across four populations, spanning intact, naturally diverged and experimentally derived genomic states. Finally, we characterized the evolutionary and sequence-compositional properties of the affected locus across *Komagataeibacter* genomes using comparative phylogenomic, including gene-tree-species-tree comparisons based on normalized Robinson-Foulds distances. Our findings reveal how routine strain handling can expose instability within evolutionarily distinct MGE-rich accessory genomic regions, generating population-level genomic variation with consequences for strain reproducibility and genome engineering.

## 2. Results

### 2.1 Serendipitous discovery of recurrent genome reduction at an IS5-flanked locus

Comparative genome analysis unexpectedly revealed a large chromosomal reduction in a *K. rhaeticus iGEM* glycerol stock prepared in 2025 (KR25), derived from a culture received as an agar-stab delivery from Ellis Lab (Imperial College London) and subsequently maintained by routine cryopreservation (see Methods). In contrast, an independently maintained 2022 stock (KR22), also received as an agar-stab, showed an intact genome across the corresponding genomic region (Figure 1).

**Figure 1.**
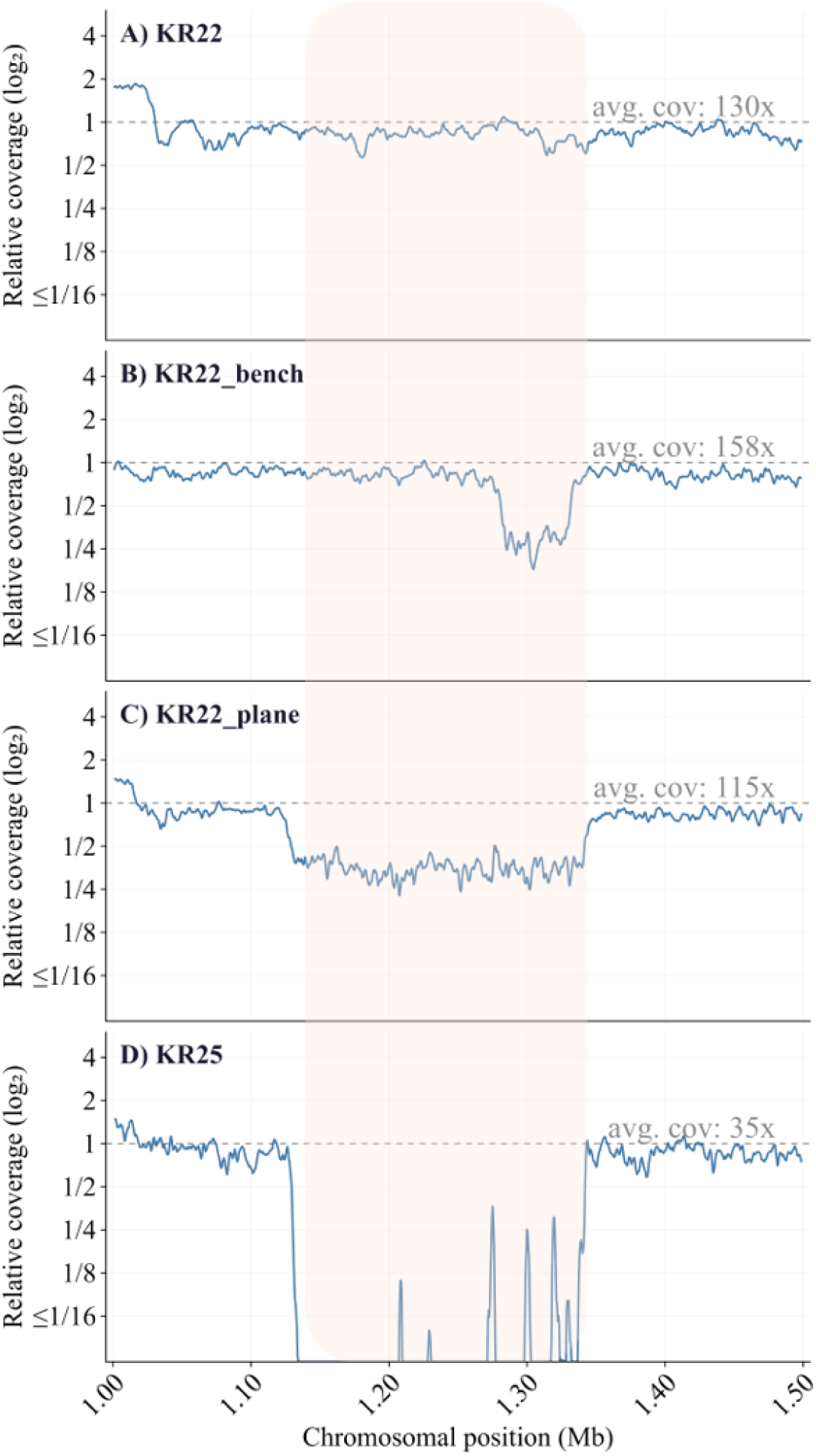
Long read sequencing coverage reveals recurrent genome reduction at a conserved chromosomal locus in *K. rhaeticus*. Relative sequencing coverage normalized to the genome-wide average (log₂ scale) is shown across the 1.00–1.50 Mb chromosomal region for the four *K. rhaeticus* populations. Average coverage (avg.cov) is indicated in each panel relative to the dashed reference line (relative coverage = 1.0). **(A)** KR22, intact reference population (avg.cov 130x). **(B)** KR22_bench, cells recovered after three weeks of agar-stab storage on a laboratory bench (avg.cov 158x). **(C)** KR22_plane, cells recovered after three weeks of agar-stab storage and air transport between Finland and Italy (avg.cov 115x). **(D)** KR25, derived from an agar-stab delivery received in 2024 (avg.cov 35x). Shading indicates the region of reduced coverage across panels B, C and D (coordinates 1,130,236–1,338,916 bp).

KR25 showed near-complete loss of sequencing coverage across a ∼210 kbp chromosomal region (coordinates 1,130,236-1,338,916 bp; NCBI BioProject PRJNA1518148), whereas KR22 retained full coverage across the corresponding locus. The region was flanked by homologous IS5-family transposase sequences, consistent with IS5-mediated homologous recombination as the mechanism for loss of the intervening sequence (1,11). Residual coverage peaks within the depleted region in KR25 were attributable to sequences homologous to IS5 elements elsewhere in the chromosome.

Because both KR22 and KR25 had been received as agar-stab cultures, the stock comparison alone could not establish whether strain handling contributed to the genome reduction. We therefore tested whether the observed genomic reduction could be reproduced under controlled agar-stab conditions using the intact KR22 population as the parental strain (see Methods). Two conditions were examined: storage of an agar-stab on a laboratory bench for three weeks (KR22_bench) and transport of an agar-stab by air between Finland and Italy over the same period (KR22_plane). Long-read sequencing revealed reduced coverage at the same chromosomal locus in both experimentally derived populations (Figure 1). KR22_plane showed depletion across the same ∼210 kbp region affected in KR25, whereas KR22_bench showed a smaller, nested deletion bounded by a distinct pair of homologous IS5 sequences. The partial coverage depletion observed in both KR22_plane and KR22_bench is consistent with the presence of deletion-bearing and intact cells within the recovered populations, indicating that genome reduction occurred at the population level rather than as a uniformly fixed clonal event.

Together, these observations identify a reproducibly unstable chromosomal region in *K. rhaeticus* that undergoes variable-scale genome reduction under agar-stab conditions, with deletion boundaries defined by homologous IS5 elements.

### 2.2 Distinct genomic states are associated with differences in growth and bacterial cellulose production

To determine whether the distinct genomic states were accompanied by phenotypic variation, we compared growth and BC production in the intact KR22 population, the evolved KR25 population, and the two experimentally derived KR22_plane and KR22_bench populations.

The populations carrying the larger genomic deletion, KR25 and KR22_plane, showed an earlier growth onset than the intact KR22 population and KR22_bench population (Figure 2). KR22_plane exited the lag phase at ∼20 h and KR25 at ∼26 h, compared with ∼48 h for KR22 and ∼44 h for KR22_bench. KR22_plane also showed the highest specific growth rate (0.0599± 0.0230), followed by KR25 (0.0550 ± 0.0080), KR22 (0.0419 ± 0.0070) and KR22_bench (0.0217 ± 0.0011) (Figure 2). The two populations carrying the larger genome reduction also reached the highest maximum backscatter values, with KR22_plane maintaining an increased signal through 336 h (Supplementary File 1: Figure S1).

**Figure 2.**
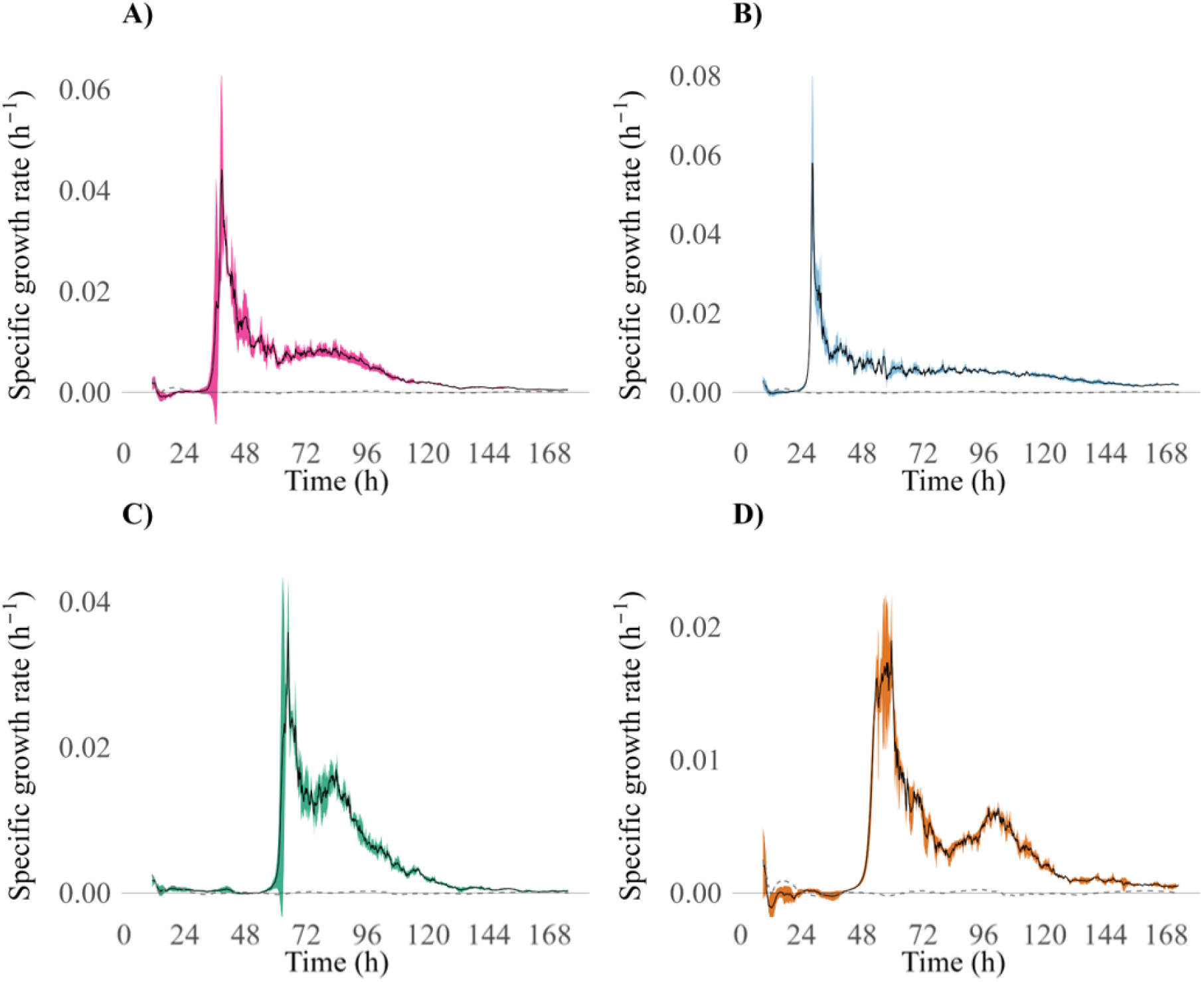
Growth rate dynamics of the four *K. rhaeticus* populations. Mean specific growth rate (h⁻¹) derived from CGQ backscatter measurements, is shown for four *K. rhaeticus* populations during shaken cultivation in HS-glucose medium supplemented with cellulase. **(A)** KR25, carrying a complete deletion of the ∼210 kbp region. **(B)** KR22_plane, a heterogeneous population carrying the same ∼210 kbp deletion. **(C)** KR22, wild-type reference with an intact genome. **(D)** KR22_bench, a heterogeneous population carrying a partial ∼50 kbp deletion nested within the larger region. Each panel shows the mean (solid line) ± standard deviation (SD; shaded band) of three biological replicates. The grey dashed line represents the mean from medium-only blank controls.

Independent growth measurements were consistent with the continuous growth profiles. At 336 h, KR22_plane reached the highest optical density (OD_600nm_; 2.38 ± 0.01), followed by KR25 (1.24 ± 0.18), KR22 (1.13 ± 0.05) and KR22_bench (1.02 ± 0.02).

We next quantified BC production after 10 days of static cultivation (Supplementary File 1: Figure S2). BC titer was highest in KR22_plane (1.76 ± 0.21 g/L), representing an 80.8% increase relative to KR22 (0.97 ± 0.10 g/L; Welch’s t-test, t = 2.84, df = 5.90, p = 0.011), followed by KR25 (1.35 ± 0.14 g/L; 42.4% increase relative to KR22; Welch’s t-test, t = 3.77, df = 4.48, p = 0.013). KR22_bench produced the lowest BC yield of all four populations (0.64 ± 0.03 g/L), representing a 33.8% decrease relative to KR22 (Welch’s t-test, t = 2.34, df = 5.57, p = 0.021). Notably, KR22_plane recapitulated the phenotypic trend observed in KR25 despite being independently derived from the intact KR22 population. Together, these observations indicate that distinct genomic states at the unstable locus can be accompanied by distinct phenotypes.

### 2.3 Phylogenomic characterization of the deleted region

To determine whether the recurrently unstable region differs in its evolutionary characteristics from the surrounding genomic background, we characterized the evolutionary distribution of the 156 protein-coding genes within the region.

We reasoned that characterizing the evolutionary history of the region, and specifically whether the gene content reflects horizontal acquisition rather than vertical inheritance would provide insights into the genomic properties underlying its instability. Annotated transposases, ISs, and invertases were excluded from the analysis. Homologous sequences were identified across a dataset of 39 bacterial genomes (Supplementary File 2), comprising 16 *Komagataeibacter/Novacetimonas* strains (20) and 23 phylogenetically diverse outgroup species. Gene presence and absence were assessed across the dataset (Figure 3A), and orthologous groups and their corresponding gene-trees were inferred using OrthoFinder (21). Gene-trees were compared with the species-tree, and the degree of phylogenetic incongruence was quantified using normalized Robinson-Foulds (RF) distance, where 0 indicates identical topology and 1 indicates maximal topological discordance (Figure 3B; see Methods).

**Figure 3.**
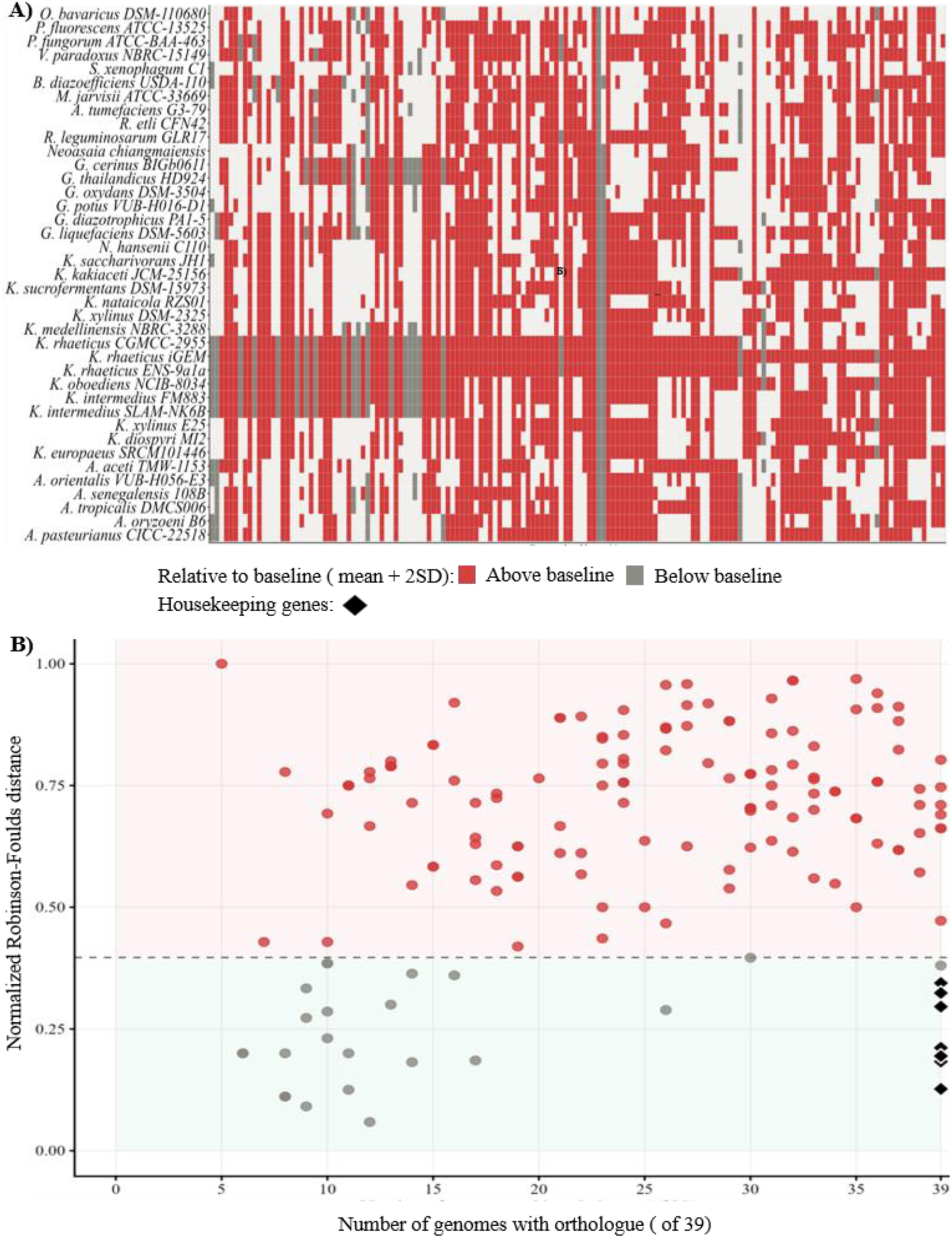
Conservation of breadth and gene-tree conflict for genes within the ∼210 kbp deleted region. **(A)** Presence/absence of the 156 genes across the 39 compared genomes. Genes are ordered by genomic position within the deleted region (x-axis) and genomes are ordered according to their phylogenetic relationships (y-axis). Brackets are coloured according to the Robinson-Foulds (RF) values of each gene, corresponding to the values shown in panel B. **(B)** Relationship between conservation breadth and phylogenetic incongruence for the 156 protein-coding genes within the deleted region (excluding transposases, ISs and invertases). Each point represents one gene; the x-axis shows the number of genomes in which an orthologue was detected (conservation breadth), and the y-axis shows the normalised RF distance between each gene-tree and species-tree. Points are coloured according to their position relative to an empirically derived baseline threshold (dashed horizontal line), calculated as the mean RF distance across eight single-copy housekeeping genes (*ligA*, *rpoA*, *rpoB*, *ruvB*, *atpA*, *polA*, *dnaK. serS*; black diamonds) + 2 SD.

The presence/absence analysis revealed substantial heterogeneity in gene conservation across the region, with individual genes ranging from broadly conserved to restricted to subsets of the 39 genomes (Figure 3A). This heterogeneity indicates that the region does not behave as a uniformly conserved genomic block and instead comprises of genes with distinct evolutionary distributions across *Komagataeibacter* and the outgroup taxa.

Gene-trees/species-tree comparison revealed widespread phylogenetic incongruence across the region (22) (Figure 3A). Of the 156 genes analysed, 132 (84.6%) had normalized RF distances above the empirical baseline threshold (baseline mean + 2 standard deviations = 0.397), whereas 24 genes (15.4%) fell below. Among the genes exceeding the threshold, 15 genes representing 14 unique orthogroups showed particularly high phylogenetic conflict (RF ≥ 0.9) together with variable conservation breadth. One gene (coordinates 1,279,468-1,280,082 bp; NCBI BioProject PRJNA1518148), encoding a nuclease-related domain-containing protein, showed maximal phylogenetic discordance (RF = 1.0). Homologues of this gene were detected only within *Komagataeibacter*, suggesting a history of horizontal gene transfer within the genus. However, because only five genomes contained homologues of this gene, its individual gene-tree topology provides limited resolution for precisely identifying its evolutionary scenario. The evolutionary heterogeneity of the region is further supported by its sequence composition and MGEs content. Twelve IS elements are distributed across this region, representing approximately twice the genome-wide mean density (12 observed versus a mean value of 5.99 per equivalent genomic window). This IS element content was predominantly represented by the IS5/IS1032 family, which flanks the deletion boundaries. Indeed, the deleted region contained eight IS5/IS1032 elements, representing a 4.5-fold enrichment relative to the genome-wide mean (p < 0.0001) with no randomly sampled 210kbp region containing an equivalent number of copies. The GC content of the deleted region ranged from 39.4% to 70.1% (mean 60.17%; Supplementary File 1: Figure S3). The combination of variable gene conservation, widespread gene-tree/species-tree incongruence, above average IS element density and heterogeneous GC content, distinguishes the region from a uniformly inherited genomic background and supports its characterization as a dynamic accessory genomic region with a heterogenous evolutionary history (1,22,23)

These evolutionary characteristics provide a genomic context for the recurrent structural instability observed at this locus, linking its heterogeneous accessory-genome composition with its propensity for large-scale sequence loss under agar-stab conditions.

## 3. Discussion

Our findings demonstrate that routine strain handling can cause structural genomic instability in *K. rhaeticus*. A recurrently unstable accessory region underwent variable-scale genome reduction under agar-stab handling conditions, with distinct genomic states associated with differences in growth and BC production. This extends previous observations of genome instability in *Komagataeibacter* under cultivation stress (16,24) and suggests that routine strain maintenance and distribution can contribute to genomic and phenotypic heterogeneity. Such strain-handling-associated genetic heterogeneity has been observed in other bacteria. Spira et al. (18) previously reported that *E. coli* K-12 cultures shared via agar-stabs became genetically heterogeneous, although the identified mutations remained confined to point mutations rather than large-scale chromosomal changes. Renda et al. (25) found that *A.baylyi* lineages evolved for over 1000 generations, all the independently accumulated different mutations, including IS1236 insertions, IS1236-mediated deletions, single nucleotide polymorphisms and small indels, alongside the deletion of a 49 kbp prophage region mediated by recombination between flanking homologous 51-bp sequences.

The identification of a recurrently unstable genomic region and its experimental reproduction from an intact parental population are central findings of this study. The difference between independently maintained stocks could not establish when the reduction had occurred, as both stocks were derived from an agar-stabbed culture. Reproducing the genome reduction from an intact population under simulated conditions demonstrates that the affected locus has a reproducible propensity for structural variation under the tested conditions. The different extents of sequence loss observed among the experimental populations further indicate that the instability at the locus is not an all-or-nothing event but can generate heterogeneous population-level genomic states.

The locus architecture provides a plausible mechanistic explanation. Homologous IS5/IS1032 elements flank the locus and coincide with the deletion boundaries, consistent with IS element-mediated recombination as a mechanism for sequence loss and with the established capacity of IS elements to drive large-scale genome rearrangements in bacteria (11,25,26). While homologous IS5/IS1032 elements similarly flank other chromosomal regions in *K. rhaeticus* iGEM, including ribosomal RNA sequences and metabolic gene clusters (coordinates bp 724,118-825,481; 1,879,038 -2,026,645), none showed evidence of structural rearrangements in any of the populations examined. In the deleted region, however, IS5/IS1032 elements are significantly enriched as shown by the permutation test.

The phylogenomic analysis provides a broader context for the recurrent instability observed at this locus. The genes within the locus showed heterogeneous conservation across *Komagataeibacter* and related bacteria, alongside widespread gene-tree/species-tree discordance. Together with the presence of several IS elements and the variable GC content across the region, these features distinguish the locus from a uniformly conserved chromosomal segment and provide a plausible evolutionary context for its recurrent structural instability (22).

What is not immediately evident is why this locus becomes unstable under agar-stab conditions. Agar storage and transport impose a combination of varying physiological states and environmental conditions that were not separated in our experiments. Therefore, the study cannot distinguish whether instability is associated with a specific environmental trigger or arises more generally from changes in growth, storage, transport, or recovery conditions. Despite this, our findings show that agar-stab handling reproducibly exposes structural variation at a specific genomic location with a distinctive genomic architecture in *K. rhaeticus*

The phenotypic observations indicate that the genomic reduction was associated with functional consequences. Populations with larger genome reductions showed a common direction of change in growth and BC production, whereas the population carrying the smaller, nested deletion showed reduced growth and BC production relative to the intact population. This argues against a simple model in which genome reduction is intrinsically beneficial. The consequences of structural variations may also depend on which portion of the accessory region is lost and on the genetic background in which the deletion occurs. However, these phenotypes cannot be attributed exclusively to the deletions based on the present data, and additional unidentified genomic or physiological differences between populations cannot be excluded. Resolving these contributions will require reconstruction of defined deletions in an otherwise isogenic background and, ideally, restoration of the deleted region.

These findings have implications for the stability and reproducibility of bacterial strains used in research and biotechnological applications. The recurrent loss of an accessory region could represent an important source of strain-to-strain variation, particularly for *Komagataeibacter* strains used in synthetic biology and industrial BC production. Phenotypic discrepancies between laboratories working with ostensibly identical strains may, in some cases, reflect undetected genomic divergence introduced during distribution and storage rather than genuine biological variation. Conversely, unstable genomic regions may indicate a genetic content that is dispensable under laboratory conditions and therefore serve as engineering targets for genome streamlining. The present findings do not establish that spontaneous deletion itself is beneficial or that this specific region should be removed for chassis optimisation. Rather, they provide a basis for targeted functional testing of individual genes or subregions to determine whether defined genome reductions can improve performance reproducibly.

More broadly, our findings suggest that bacterial strain maintenance and sharing are not necessarily genomically neutral. Accessory genomes can retain evolutionary and structural dynamics and routine handling can expose these dynamics as population heterogeneity and phenotypic variation, creates a dual challenge. Accessory-genome instability may undermine reproducibility, while simultaneously revealing opportunities to identify genomic regions that influence strain performance. Identifying the genomic features that predispose accessory regions to instability may therefore provide a strategy to improve strain reproducibility and uncover new opportunities for rational genome engineering.

## 4. Conclusions

This study demonstrates that routine agar-stab handling can trigger recurrent genome rearrangement at a specific locus in *K. rhaeticus* iGEM. The affected locus exhibited distinctive phylogenomic and sequence-compositional characteristics, providing a plausible genomic context for its structural instability. Our findings have two complementary implications. First, routine strain handling practices can be a source of genotypic and phenotypic variation, particularly in organisms with MGE-rich genomes, with consequences for reproducibility in biotechnological applications. Systematic genomic screening of *Komagataeibacter* stocks following agar-stab delivery or prolonged storage could therefore help identify genomic changes that might otherwise remain undetected despite their phenotypic consequences. This consideration may also extend to other bacteria with dynamic accessory genomes. Second, identifying recurrently unstable accessory regions provides candidates for targeted investigation of genomic content that may be dispensable under defined laboratory conditions. Extending the phylogenomic analysis applied here to the full genome and to organisms with similarly dynamic accessory content, could enable systematic identification of regions with atypical evolutionary histories, offering a genome-wide perspective on genomic instability and a rational basis for targeted genome streamlining efforts. However, instability alone does not establish dispensability or suitability for genome streamlining. Defined deletion and complementation experiments will be required to determine whether specific genome reductions can reproducibly improve strain performance. Such studies could ultimately inform strategies to improve BC production and maintain genetically stable *Komagataeibacter* chassis for industrial biotechnology.

## 5. Methods

### 5.1 *K. rhaeticus* iGEM populations and cultivations

#### 5.1.1 Growth media

All cultivations were conducted in Hestrin-Schramm (HS)-glucose medium (16). For shaken cultivations, the medium was supplemented with 2% (v/v) of liquid cellulase from *T. reesei* (Biozol, Germany) to prevent cellulose accumulation and ensure biomass measurements reflect cell density.

#### 5.1.2 Strains and agar-stab delivery simulation

KR22 and KR25 were independently derived from *K. rhaeticus* iGEM cultures (27) received as agar-stab cultures from Ellis Lab (Imperial College London). These cultures were subsequently maintained through routine culture preservation and cryopreservation methods.

To simulate an agar-stab delivery, 50 µL of KR22 glycerol stock were inoculated into a 2 mL microcentrifuge tube containing 1.5 mL of HS-glucose medium supplemented with 1.5% (w/v) bacteriological agar. Both tubes were sealed with Parafilm inside a zip-lock bag. KR22_bench was stored on a laboratory bench at Aalto University (Finland) for three weeks. KR22_plane was transported by air between Finland and Italy and back over the same period, with agar-stab maintained at room temperature throughout the transport. After three-weeks, the cells were recovered by transferring the cells from the solidified medium into 10 mL of HS-glucose medium and incubating statically at 30°C for 5 days to allow BC pellicle formation. The pellicles were lysed overnight using 2% (v/v) cellulase to release the entrapped cells and subsequently washed three times with sterile 1× phosphate-buffered saline (PBS). Washed cells were then used for genomic DNA extraction and preserved as glycerol stocks (28).

#### 5.1.3 Static and shaken cultivations

For preculture preparation, 100 µL of glycerol stock was inoculated into 10 mL of HS-glucose medium and incubated statically at 30°C for 5 days. BC was lysed and cells were washed three times with 1X PBS prior to inoculation(16). BC production was assessed in 24-well plates (Axygen Scientific Inc., Union City, CA, USA) containing 8 mL of HS-glucose medium at 30°C for 10 days, in biological triplicates alongside three uninoculated controls. All cultures were initiated at an OD_600nm_ of 0.1. BC pellicles were harvested and processed as previously described (3). Statistical comparison of BC production between populations was performed using Welch’s two-sample-t-test, with comparisons made against the intact genome reference population, KR22.

Growth kinetics were assessed under shaken conditions (30°C, 220 rpm for 14 days) in 250 mL Erlenmeyer flasks containing 50 mL of HS-glucose medium supplemented with 2% (v/v) cellulase. Cultures were initiated at an OD_600nm_ of 0.1 and were conducted in biological triplicates alongside two non-inoculated controls. Growth dynamics was monitored non-invasively using a Cell Growth Quantifier (CGQ; Aquila Biolabs, Scientific Bioprocessing). Backscattering data were recorded using the DOTS software, which was used to derive the specific growth rates. At the end of the cultivation, 2 mL were sampled from each flask to measure OD_600nm_ values.

Growth rates were normalised to the mean of the first 10 measurements points for each flask. Backscatter plots curves and growth-rate profiles were processed and visualized in RStudio (v2026.7.1.147) using the tidyverse packages, dplyr, tidyr, purrr and stringr (29). Figures were generated using the R packages ggplot2 (30), extrafont (31) and patchwork (32) (Supplementary File 4).

### 5.2 Genomic DNA extraction, sequencing and phylogenomic analysis

Genomic DNA was extracted as previously described (16). The genomic DNA from the analysed *K. rhaeticus* populations, were sequenced using Oxford Nanopore GridION sequencer and libraries were prepared using the Oxford Nanopore Technologies SQK-NBD114.24 barcoding kit, according to the manufacturer’s instructions (ONT, Oxford, UK). Base calling was performed using Dorado (v 7.13.6) in high-accuracy mode with barcode trimming enabled.

#### 5.2.1 Whole genome alignment and deletion detection

Reads were aligned against a *K. rhaeticus* iGEM hybrid assembly (NCBI BioProject PRJNA1518148), generated using Illumina paired-end (2×150 bp) and Oxford Nanopore (R10.4.1) (Plasmidsaurus, UK) sequencing, using Geneious Prime (v2025.2.2, Geneious Mapper). Per-base sequencing coverage was exported and processed in RStudio (v2026.7.1.147). Genome-wide average coverage was calculated as the arithmetic mean of per-base coverage while the coverage across the ∼210 kbp locus was calculated using a sliding window approach (window size: 2,000 bp, step size: 500 bp), normalised to the genome-wide average, and visualised on a log₂ scale using the RStudio ggplot2 (30), extrafont (31) patchwork (32) and scales (33) packages (Supplementary File 4).

#### 5.2.2 Phylogenomic characterisation of the affected locus

To construct a reference dataset for phylogenomic analysis, 39 bacterial genomes were selected as follows. *Komagataeibacter* and *Novacetimonas* genomes were manually curated prioritizing genome completeness and coverage. Outgroups were selected from BlastP searches using Bakta (v1.12.0) annotated genes within the ∼210 kbp deleted region as queries (34,35). Candidate species were ranked by the number of matching genes, and one to two representative genomes were selected from the top-ranking lineages. For both *Komagataeibacter*/*Novacetimonas* and outgroup genomes, assembly completeness was assessed using BUSCO (v6.1.0) (36) against appropriate lineage-specific orthologue databases and only genomes meeting a threshold of >95% were retained. This approach yielded genomes that spanned *Acetobacteraceae (Komagataeibacter, Novacetimonas, Acetobacter*, *Gluconobacter*, *Gluconacetobacter* and *Neoasaia* species) as well as phylogenetically distant lineages including *Bradyrhizobium*, *Pseudomonas*, *Mesorhizobium*, *Rhizobium*, *Paraburkholderia*, *Variovorax*, *Agrobacterium* and *Occallatibacter*. Together with the *K. rhaeticus* genome, this yielded a final dataset of 39 genomes (Supplementary File 2). Orthologous groups and gene-trees were inferred using OrthoFinder (v2.5.5) (21) with its default options. The species-tree was reconstructed de novo using GTDB-Tk (v2.7.2) (37), with *de_novo_wf* workflow and the bac120 marker gene set. All steps involving BlastP, Bakta, BUSCO, OrthoFinder, GTDB-Tk and output parsing were implemented using Python-based workflows with the corresponding script available in a public GitHub repository (https://github.com/vimagora/genome_reduction_komagataeibacter).

Gene-trees corresponding to the annotated genes within the deleted region, excluding ISs, transposases and invertases, were extracted from the OrthoFinder output, yielding 147 gene-trees for comparison against the species-tree. As the Robinson-Foulds (RF) distance is defined for trees with uniquely labelled tips (39), gene-trees containing multiple tips from the same species due to intra-genome paralogy were reduced to a single representative tip per taxon, avoiding the complications associated with repeated species labels in multi-copy gene-trees (40). The representative tip was selected arbitrarily, with no intent to resolve paralogous relationships. Consequently, the resulting RF distances were interpreted as species-level topology comparisons rather than analyses of paralogous evolutionary histories. Following the removal of a small number of genes with insufficient taxon representation or transposase-associated annotations not identified during the initial curation, a final set of 156 genes were used for RF distance analysis (Supplementary File 3). Phylogenetic incongruence between each gene-tree and species-tree was quantified using the normalised RF distance (39), calculated using the RF.dist function (with normalize = TRUE) in the Phangorn package (v2.12.1) (41) in RStudio. Prior to RF distance calculation, each gene-tree and species-tree were pruned to their shared taxon set. Conservation breadth for each gene was defined as the number of the 39 genomes in which an orthologue was detected by OrthoFinder. To establish an empirical reference baseline of phylogenetic discordance, RF distances were additionally computed for eight single-copy orthologous housekeeping genes, *ligA*, *rpoA*, *rpoB*, *ruvB*, *atpA*, *polA*, *dnaK* and *serS* (6,42–44). A gene-tree distance from species-tree was considered significantly incongruent if the RF distance exceeded the baseline mean by more than two standard deviations (baseline mean = 0.226, SD = 0.086; threshold = 0.397) (45). Although housekeeping genes set (n = 8) is consistent with conventions in multilocus sequence analysis (46), this threshold should be interpreted as an approximate, literature-motivated benchmark rather than a precise statistical boundary. All the RStudio scripts implementing the RF analyses, baseline calculations and associated visualizations are provided in Supplementary File 4 and relied R packages hereby cited (29–32,41,47,48).

To assess IS element enrichment within the deleted region, the locus was assessed using a permutation test. Prior to the permutation test, the Bakta annotation was extended to the full chromosome. IS elements were defined as annotated CDS whose product description specifically contained an IS family designation. IS21 elements, which encode two separately annotated subunits (IstA and IstB), were collapsed into a single count. A total of 10,000 non-overlapping windows (210 kbp in size) were randomly sampled from the chromosome, excluding windows overlapping the deleted region. The permutation analysis was implemented in Python with the corresponding scripts available on a public GitHub repository (38).

GC content across the deleted region was assessed using a sliding-window approach in RStudio, with window size of 1 kbp and a step size of 500 bp. Visualisation was performed using ggplot2 and extrafont packages (30,31) and the corresponding RStudio script is provided in Supplementary File 4.

## 5.3 Data availability

The *K. rhaeticus* iGEM hybrid assembly used as a reference genome has been submitted to NCBI (BioProject PRJNA1518148). Outgroup and *Komagataeibacter* genomes used in phylogenomic analyses are publicly available on NCBI, with their GenBank accession numbers, assembly statistics and BUSCO completeness score provided in Supplementary File 2. Supplementary File 3 lists annotated genes within the deleted region, excluding transposases, ISs and invertases, alongside their ortholog matches and RF distances. The RStudio scripts can be found as Supplementary File 4. Python scripts implementing BlastP searches, Orthofinder, BUSCO, GTDB-tk and the IS permutation test can be consulted on GitHub (38).Raw Nanopore FASTQ files for KR22, KR25, KR22_bench and KR22_plane, will be available on Zenodo (DOI:10.5281/zenodo.22895627) or upon request.

## Acknowledgments

D.B. and R.M. acknowledge the Center for Young Synbio Scientists for supporting this study. R.M. would like to acknowledge the Research Council of Finland (Grant Numbers 353673 and 372125) for supporting this study. M.R.M. acknowledges funding from the Research Council of Finland (Grant numbers: 348443 and 366183) and the Novo Nordisk Foundation (Grant number: NNF21OC0067087). D.B. further acknowledges CSC–IT Center for Science, Finland, for providing the computational resources. The authors thank Assoc. Prof Paula Jouhten and Dr. An Nguyen for assisting with the CGQ and Dr Gianluca Prezza for uploading the *K. rhaeticus* iGEM hybrid assembly to NCBI.

## Authorś contribution

D.B. designed and performed the shaking cultivation with CGQ, conducted the bioinformatics analyses, including comparative genome analysis and phylogenomic analysis; performed the sequencing; interpreted the data and drafted the original manuscript. V.M.G.R. supported the phylogenomic analysis, and gene-tree and species-tree construction, and M.R.M. supervised V.M.G.R. S.H. conducted the initial *K. rhaeticus* stock screening and BC production tests. A.P. conducted the hybrid sequencing of the bioinformatic reference genome used in this study and T.E. supervised A.P. R.M. conceptualized the project, interpreted the data, revised the manuscript, and supervised the work. All authors contributed to manuscript revision.

## Large Language Models use

Large Language Models assisted with RStudio packages implementation and script development. All scripts were manually reviewed and executed in RStudio.

## References

1. Siguier P, Gourbeyre E, Chandler M. Bacterial insertion sequences: their genomic impact and diversity. FEMS Microbiol Rev. 2014 Sep;38(5):865–91. doi:10.1111/1574-6976.12067

2. Malcı K, Li IS, Kisseroudis N, Ellis T. Modulating Microbial Materials - Engineering Bacterial Cellulose with Synthetic Biology. ACS Synth Biol. 2024 Dec 20;13(12):3857–75. doi:10.1021/acssynbio.4c00615

3. Mangayil R, Rajala S, Pammo A, Sarlin E, Luo J, Santala V, et al. Engineering and Characterization of Bacterial Nanocellulose Films as Low Cost and Flexible Sensor Material. ACS Appl Mater Interfaces. 2017 Jun 7;9(22):19048–56. doi:10.1021/acsami.7b04927

4. Vannas J, Singh A, Hannikainen B, De Assis MA, Ledesma-Amaro R, Ellis T, et al. A signal peptide-guided approach towards in situ functionalization of bacterial nanocellulose in Komagataeibacter rhaeticus. J Biol Eng. 2026 Jul 17. doi:10.1186/s13036-026-00734-w

5. Walker KT, Li IS, Keane J, Goosens VJ, Song W, Lee KY, et al. Self-pigmenting textiles grown from cellulose-producing bacteria with engineered tyrosinase expression. Nat Biotechnol. 2025 Mar;43(3):345–54. doi:10.1038/s41587-024-02194-3

6. Mangayil R, Sarlin E, Ellis T, Santala V. Modulating bacterial nanocellulose crystallinity through post-transcriptional repression in Komagataeibacter xylinus. Carbohydr Polym Technol Appl. 2025 Mar;9:100734. doi:10.1016/j.carpta.2025.100734

7. Deng S, Wang L, Chen G, Qin Q, Dong S, Zhang H. Complete genome analysis of the cellulose producing strain Komagataeibacter sucrofermentans SMEG01. Sci Rep. 2025 Jul 2;15(1):23102. doi:10.1038/s41598-025-07045-y

8. Akhoon BA, Qiao Q, Stewart A, Chen J, Rodriguez Lopez CM, Corbin KR. Pangenomic analysis of the bacterial cellulose-producing genera Komagataeibacter and Novacetimonas. Int J Biol Macromol. 2025 Apr;298:139980. doi:10.1016/j.ijbiomac.2025.139980

9. Ryngajłło M, Kubiak K, Jędrzejczak-Krzepkowska M, Jacek P, Bielecki S. Comparative genomics of the *Komagataeibacter* strains—Efficient bionanocellulose producers. MicrobiologyOpen. 2019 May;8(5):e00731. doi:10.1002/mbo3.731

10. Frost LS, Leplae R, Summers AO, Toussaint A. Mobile genetic elements: the agents of open source evolution. Nat Rev Microbiol. 2005 Sep;3(9):722–32. doi:10.1038/nrmicro1235

11. Lee H, Doak TG, Popodi E, Foster PL, Tang H. Insertion sequence-caused large-scale rearrangements in the genome of *Escherichia coli*. Nucleic Acids Res. 2016 Jul 18;gkw647. doi:10.1093/nar/gkw647

12. Darmon E, Leach DRF. Bacterial Genome Instability. Microbiol Mol Biol Rev. 2014 Mar;78(1):1–39. doi:10.1128/MMBR.00035-13

13. Schramm M, Hestrin S. Factors affecting Production of Cellulose at the Air/ Liquid Interface of a Culture of Acetobacter xylinum. J Gen Microbiol. 1954 Aug 1;11(1):123–9. doi:10.1099/00221287-11-1-123

14. Hur DH, Rhee HS, Lee JH, Shim WY, Kim TY, Lee SY, et al. Enhanced production of cellulose in Komagataeibacter xylinus by preventing insertion of IS element into cellulose synthesis gene. Biochem Eng J. 2020 Apr;156:107527. doi:10.1016/j.bej.2020.107527

15. Coucheron DH. An Acetobacter xylinum insertion sequence element associated with inactivation of cellulose production. J Bacteriol. 1991 Sep;173(18):5723–31. doi:10.1128/jb.173.18.5723-5731.1991

16. Bersanetti D, Mangayil R. Exploring Genome-Wide Mutation Dynamics and Bacterial Cellulose Impairment in *Komagataeibacter intermedius* Cultivated Under Agitation Stress. Comput Struct Biotechnol J. 2026 Mar 1;35(1):0009. doi:10.34133/csbj.0009

17. Sakurai K, Kawasaki H. Genetic variation during long-term preservation of bacteria in public culture collections. Int J Syst Evol Microbiol. 2018 May 1;68(5):1815–21. doi:10.1099/ijsem.0.002717

18. Spira B, De Almeida Toledo R, Maharjan RP, Ferenci T. The uncertain consequences of transferring bacterial strains between laboratories - rpoS instability as an example. BMC Microbiol. 2011;11(1):248. doi:10.1186/1471-2180-11-248

19. Kanai Y, Shibai A, Yokoi N, Tsuru S, Furusawa C. Laboratory evolution of the bacterial genome structure through insertion sequence activation. Nucleic Acids Res. 2025 May 10;53(9):gkaf331. doi:10.1093/nar/gkaf331

20. Brandão PR, Crespo MTB, Nascimento FX. Phylogenomic and comparative analyses support the reclassification of several Komagataeibacter species as novel members of the Novacetimonas gen. nov. and bring new insights into the evolution of cellulose synthase genes. Int J Syst Evol Microbiol. 2022 Feb 17;72(2). doi:10.1099/ijsem.0.005252

21. Emms DM, Kelly S. OrthoFinder: phylogenetic orthology inference for comparative genomics. Genome Biol. 2019 Dec;20(1):238. doi:10.1186/s13059-019-1832-y

22. Ravenhall M, Škunca N, Lassalle F, Dessimoz C. Inferring Horizontal Gene Transfer. Wodak S, editor. PLOS Comput Biol. 2015 May 28;11(5):e1004095. doi:10.1371/journal.pcbi.1004095

23. Zhang R, Ou HY, Gao F, Luo H. Identification of Horizontally-transferred Genomic Islands and Genome Segmentation Points by Using the GC Profile Method. Curr Genomics. 2014 Apr 31;15(2):113–21. doi:10.2174/1389202915999140328163125

24. Zhang H, Chen C, Yang J, Sun B, Lin J, Sun D. Effect of Culture Conditions on Cellulose Production by a *Komagataeibacter xylinus* Strain. Macromol Biosci. 2022 Jun;22(6). doi:10.1002/mabi.202100476

25. Renda BA, Dasgupta A, Leon D, Barrick JE. Genome Instability Mediates the Loss of Key Traits by Acinetobacter baylyi ADP1 during Laboratory Evolution. Zhulin IB, editor. J Bacteriol. 2015 Mar;197(5):872–81. doi:10.1128/JB.02263-14

26. Raeside C, Gaffé J, Deatherage DE, Tenaillon O, Briska AM, Ptashkin RN, et al. Large Chromosomal Rearrangements during a Long-Term Evolution Experiment with Escherichia coli. Molin S, Baquero F, editors. mBio. 2014 Oct 31;5(5):e01377–14. doi:10.1128/mBio.01377-14

27. Florea M, Hagemann H, Santosa G, Abbott J, Micklem CN, Spencer-Milnes X, et al. Engineering control of bacterial cellulose production using a genetic toolkit and a new cellulose-producing strain. Proc Natl Acad Sci. 2016 Jun 14;113(24). doi:10.1073/pnas.1522985113

28. Cannazza P, Rissanen AJ, Sarlin E, Guizelini D, Minardi C, Losoi P, et al. Characterization, genome analysis and genetic tractability studies of a new nanocellulose producing Komagataeibacter intermedius isolate. Sci Rep. 2022 Nov 28;12(1). doi:10.1038/s41598-022-24735-z

29. Wickham H, Averick M, Bryan J, Chang W, McGowan L, François R, et al. Welcome to the Tidyverse. J Open Source Softw. 2019 Nov 21;4(43):1686. doi:10.21105/joss.01686

30. Wickham H. ggplot2 [Internet]. Cham: Springer International Publishing; 2016. (Use R!). Available from: http://link.springer.com/10.1007/978-3-319-24277-4 doi:10.1007/978-3-319-24277-4

31. Chang W. extrafont: Tools for Using Fonts. 2012. p. 0.20. Available from: https://CRAN.R-project.org/package=extrafont doi:10.32614/CRAN.package.extrafont

32. Thomas Lin Pedersen. patchwork: The Composer of Plots [Internet]. 2025. Available from: https://patchwork.data-imaginist.com

33. Hadley Wickham, Thomas Lin Pedersen, Dana Seidel. scales: Scale Functions for Visualization. 2025. Available from: https://scales.r-lib.org

34. Schwengers O, Jelonek L, Dieckmann MA, Beyvers S, Blom J, Goesmann A. Bakta: rapid and standardized annotation of bacterial genomes via alignment-free sequence identification: Find out more about Bakta, the motivation, challenges and applications, here. Microb Genomics. 2021 Nov 30;7(11). doi:10.1099/mgen.0.000685

35. Camacho C, Coulouris G, Avagyan V, Ma N, Papadopoulos J, Bealer K, et al. BLAST+: architecture and applications. BMC Bioinformatics. 2009 Dec;10(1):421. doi:10.1186/1471-2105-10-421

36. Tegenfeldt F, Kuznetsov D, Manni M, Berkeley M, Zdobnov EM, Kriventseva EV. OrthoDB and BUSCO update: annotation of orthologs with wider sampling of genomes. Nucleic Acids Res. 2025 Jan 6;53(D1):D516–22. doi:10.1093/nar/gkae987

37. Chaumeil PA, Mussig AJ, Hugenholtz P, Parks DH. GTDB-Tk v2: memory friendly classification with the genome taxonomy database. Borgwardt K, editor. Bioinformatics. 2022 Nov 30;38(23):5315–6. doi:10.1093/bioinformatics/btac672

38. https://github.com/vimagora/genome_reduction_komagataeibacter

39. Robinson DF, Foulds LR. Comparison of phylogenetic trees. Math Biosci. 1981 Feb;53(1–2):131–47. doi:10.1016/0025-5564(81)90043-2

40. Chaudhary R, Burleigh JG, Fernández-Baca D. Inferring species trees from incongruent multi-copy gene trees using the Robinson-Foulds distance. Algorithms Mol Biol. 2013 Dec;8(1):28. doi:10.1186/1748-7188-8-28

41. Schliep KP. phangorn: phylogenetic analysis in R. Bioinformatics. 2011 Feb 15;27(4):592–3. doi:10.1093/bioinformatics/btq706

42. Sheoran N, Valiya Nadakkakath A, Munjal V, Kundu A, Subaharan K, Venugopal V, et al. Genetic analysis of plant endophytic Pseudomonas putida BP25 and chemo-profiling of its antimicrobial volatile organic compounds. Microbiol Res. 2015 Apr;173:66–78. doi:10.1016/j.micres.2015.02.001

43. Liu L xiang, Liu S xin, Wang Y mei, Bi J cai, Chen H mei, Deng J, et al. Komagataeibacter cocois sp. nov., a novel cellulose-producing strain isolated from coconut milk. Int J Syst Evol Microbiol. 2018 Oct 1;68(10):3125–31. doi:10.1099/ijsem.0.002947

44. Cai H, Rodriguez BT, Zhang W, Broadbent JR, Steele JL. Genotypic and phenotypic characterization of Lactobacillus casei strains isolated from different ecological niches suggests frequent recombination and niche specificity. Microbiology. 2007 Aug 1;153(8):2655–65. doi:10.1099/mic.0.2007/006452-0

45. Poptsova MS, Gogarten JP. The power of phylogenetic approaches to detect horizontally transferred genes. BMC Evol Biol. 2007;7(1):45. doi:10.1186/1471-2148-7-45

46. Glaeser SP, Kämpfer P. Multilocus sequence analysis (MLSA) in prokaryotic taxonomy. Syst Appl Microbiol. 2015 Jun;38(4):237–45. doi:10.1016/j.syapm.2015.03.007

47. Revell LJ. phytools 2.0: an updated R ecosystem for phylogenetic comparative methods (and other things). PeerJ. 2024 Jan 5;12:e16505. doi:10.7717/peerj.16505

48. Paradis E, Schliep K. ape 5.0: an environment for modern phylogenetics and evolutionary analyses in R. Schwartz R, editor. Bioinformatics. 2019 Feb 1;35(3):526–8. doi:10.1093/bioinformatics/bty633

